# Runs of homozygosity in killer whale genomes provide a global record of demographic histories

**DOI:** 10.1101/2020.04.08.031344

**Authors:** Rebecca Hooper, Laurent Excoffier, Karin A. Forney, M. Thomas P. Gilbert, Michael D. Martin, Phillip A. Morin, Jochen B.W. Wolf, Andrew D. Foote

## Abstract

Runs of homozygosity (ROH) occur when offspring receive the same ancestral haplotype from both parents, and, accordingly, reduce individual heterozygosity. Their distribution throughout the genome contains information on the probability of inbreeding mediated by mating system and population demography. Here, we investigate variation in killer whale demographic history as reflected in genome-wide heterozygosity, using a global dataset of 26 genomes. We find an overall pattern of lower heterozygosity in genomes sampled at high latitudes, with hundreds of short ROH (< 1Mbp) reflecting high background relatedness due to coalescence of haplotypes during bottlenecks associated with founder events during post-glacial range expansions. Across most of the species’ range, intermediate length ROH (1-10Mb) revealed long-term inbreeding in 22 of the 26 sampled killer whale genomes, consistent with the high social philopatry observed in all populations studied to date. Inbreeding coefficients (F_ROH_) were comparable to those reported in other taxa with long-term low population size, such as bonobos and the Native American Karitiana of the Brazilian Amazon. The extreme outlier in this dataset, a Scottish killer whale, was homozygous over one-third of the autosomes (41.6%) with a distinct distribution of ROH length, indicating generations of inbreeding. This exceeds autozygosity in emblematic examples of long-term inbreeding, such as the Altai Neanderthal, and eastern lowland and mountain gorillas. The fate of this Scottish killer whale population, in which no calves have been born in over two decades, may be inextricably linked to its demographic history and consequential inbreeding depression.

---

Species ranges change in response to environmental oscillation [1,2], and rapid shifts can currently be observed during ongoing global warming [3,4]. Understanding how range shifts influence the genetic diversity of natural populations at the range edge is an increasingly important conservation consideration [5]. Killer whales (*Orcinus orca*) are comparable to humans in their global distribution, having colonised all the major oceans [6]. Killer whale occurrence is correlated with ocean productivity; highest densities are therefore at high latitudes, decreasing by 1-2 orders of magnitude from the Arctic and Antarctic to the tropics (Figure S1) [6]. The Northern and Southern extremes of this range were covered by ice sheets during the Last Glacial Maximum (Figure 1) and must therefore have been colonised through range expansion from lower latitudes in the last 16 Kyr [7-9]. Independent post-glacial range expansions in different ocean basins offer the possibility to explore the genetic outcomes of independent, parallel demographic histories. Killer whales thus represent a useful system for studying the relationship between demographic history, range expansions, and genetic diversity. Here, we report the analyses of whole genome sequences from a global dataset of killer whales (Figure 1).

**Figure 1.**
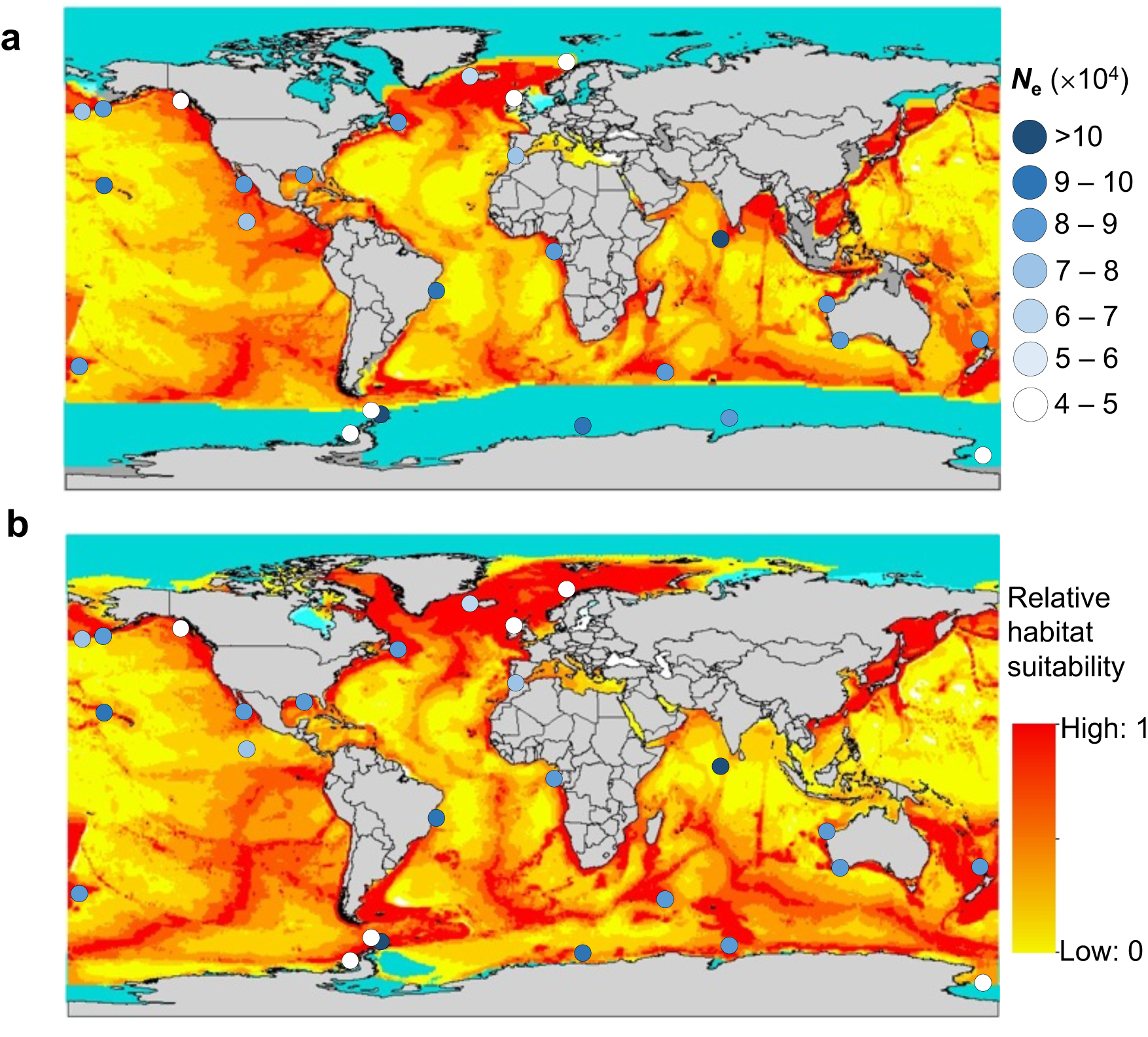
Sampling locations of the individuals included in this study overlain on a modelled suitable habitat map for killer whales. Sample markers are coloured by effective population size (*N*_e_) derived from estimates of theta (θ = 4 *N*_e_*μ*), see Supplementary Table 1. Background colours indicate **a**, hindcast prediction of core suitable habitat during the Last Glacial Maximum (LGM). **b**, core suitable habitat in the present day. Turquoise colour represents areas with >50% sea ice concentration during both time periods. Land exposed due to sea level changes during the LGM is shown in dark grey.

## RESULTS

### Heterozygosity Correlates with Latitude

To understand the impact of past range expansions and founder effects on genetic diversity in high latitude killer whale populations, we therefore sought to estimate variation in heterozygosity among our global dataset of individuals. We estimated heterozygosity of autosomal regions from individual diploid genomes corrected for coverage. For small values and under the infinite sites model [9], individual heterozygosity is a good, unbiased estimator of the population mutation rate theta (θ) [10]. Consistent with the prediction that range expansions may reduce genetic diversity [5], θ was significantly correlated with degrees latitude from the equator (Spearman correlation ρ = −0.44, P < 0.05; Figure S2) and the lowest estimates of θ were found in high latitude populations, *e*.*g*. Alaskan *resident* ecotype, Antarctic types B1, B2 and C, Iceland, Norway and Scotland (Figure 1; Table S1). However, the correlation between θ and latitude was weak due to a wide spread of values of θ at latitudes > 50° from the equator, suggesting latitude is not the only variable influencing heterozygosity. Using a general linear model (GLM; Table S2) to explore whether latitude and/or ecotype influence heterozygosity, partial R-squared values were extracted to explore the variation in heterozygosity explained by each predictor variable when controlling for the other. Results showed a significant relationship between ecotype and θ (df = 2, p < 0.001; partial R^2^ = 0.82), while correlation of latitude and θ was not significant (df = 1, p = 0.09; partial R^2^ = 0.12).

Estimates of effective population size (*N*_e_) derived from our estimates of θ (Figure 1) were concordant with previous estimates of ancestral *N*_e_ derived from IMa [12] analysis on pairs of populations using 16 microsatellite loci, which Hoelzel et al. [7] predicted could represent global *N*_e_. Low levels of gene flow among demes that retain their relative sizes can equilibrate the mean allele frequency across demes [13]. Thus, the allele frequencies, coalescence times and genetic diversity of neutral loci within a single deme are expected to reflect that of a stationary panmictic population with *N*_e_ equal to the sum of all individual demes in a structured population [13]. Whilst killer whale populations may violate some of the underlying assumptions of these expectations, low levels of migration among populations in low density regions appear to retain genetic diversity at low latitudes.

To confirm that variation in heterozygosity among our 26 genome sequences accurately represented global patterns, we analysed a published dataset of 91 SNP genotypes from 128 samples sequenced to a depth of 11.6× coverage [8]. Heterozygosity was significantly correlated with latitude in both the Northern (Spearman correlation ρ = −0.30, P < 0.01) and Southern (Spearman correlation ρ = 0.32, P < 0.05) hemispheres (Figure S3). However, again many of the data points were not a good fit to the regression line (Figure S3), suggesting variables other than latitude may also drive variation in heterozygosity. A GLM with both ecotype and latitude as predictor variables showed that among North Pacific samples both latitude and ecotype explained a significant proportion of variance in heterozygosity (latitude: df = 1, *p* = 0.01; ecotype: df = 3, *p* = <0.001), with ecotype explaining more variance in heterozygosity than latitude (partial R^2^ for ecotype = 0.37; partial R^2^for latitude = 0.09). For example, the *resident* fish-eating ecotype had significantly lower observed heterozygosity than the sympatric *transient* mammal-eating ecotype, despite occupying overlapping latitudes in the North Pacific (Table S3, Figure S4). Similarly, in the Southern Ocean, ecotype significantly explained variation in heterozygosity (df = 3, p <0.001, partial R^2^ = 0.77), but latitude did not (df = 1, p = 0.39, partial R^2^ = 0.02). The Antarctic types (B1, B2 and C), which are found around the Antarctic continent and share a recent demographic history [8,9,14], had significantly lower heterozygosity than other Southern Ocean individuals, but did not significantly differ from each other (Table S4, Figure S5). Thus, the individual demographic histories of different populations are more important at explaining variation in heterozygosity than the latitude at which populations were sampled.

### Runs of Homozygosity Reflect Demographic History

Runs of homozygosity (ROH) occur when offspring receive the same ancestral haplotype from both parents due to inbreeding [15]. The number and length of ROH reflect the timing and intensity of a population bottleneck, with longer ROH originating from recent inbreeding, and shorter ROH from ancestral bottlenecks [15]. We therefore called ROH in individuals from our global dataset using the sliding window approach implemented in PLINK [16] to compare among populations and gain insights into how population history could explain the observed patterns of ROH (Figure 2).

**Figure 2.**
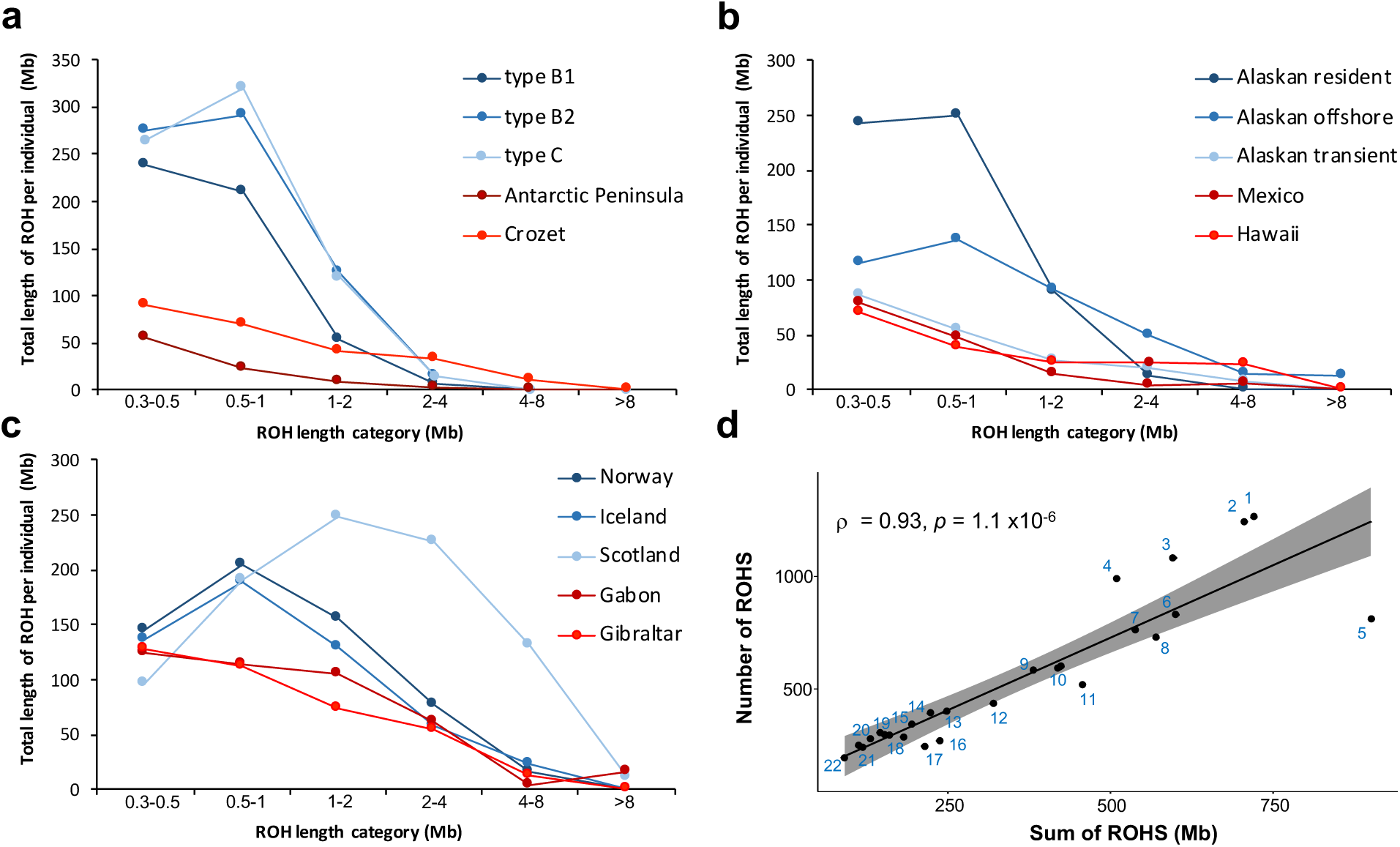
Sum of runs of homozygosity (ROH) in different length categories. Genomes from **a**, the North Pacific, **b**, the Southern Ocean and Antarctica, and **c**, the North Atlantic. **d**, Number of ROHs compared to the total length of ROHs across the autosome. Numbered data points indicate the following samples: 1. Antarctic *type C*, 2. Antarctic *type B2*, 3. Alaskan *resident*, 4. Antarctic *type B1*, 5. Scotland, 6. Norway, 7. Iceland, 8. ETP (Clipperton Island), 9. Gibraltar, 10. Gabon and Alaskan *offshore*, 11. New Zealand, 12. Newfoundland, 13. Crozet, 14. Chatham Islands, 15. Alaskan *transient*, 16. Gulf of Mexico, 17. Brazil, 18. Hawaii, 19. W. Australia, ETP (Mexico) and Southern Ocean, 20. Maldives, 21. SW. Australia and Southern Ocean, 22. Antarctic Peninsula. Shading indicates the 95% confidence intervals of the forecasted regression line.

The sum and number of ROH within individual genomes were highly correlated (Spearman correlation ρ = 0.93, P = 1.1 ×10^−6^; Figure 2D). Qualitatively similar ROH length distributions were observed among individuals from high latitude populations in different ocean basins (Figure 2A-C). Autozygosity was spread across hundreds of short (less than 1Mb in length) ROH in high latitude populations from the North Atlantic, Antarctic and North Pacific. The sum of short ROH across each individual genome was positively and significantly correlated (Spearman correlation ρ = 0.40, P < 0.05) with degrees latitude from the equator (Figure S6A). A GLM (Table S5) with the sum of short ROH as the response variable, and both latitude and ecotype as predictors, found a significant effect of ecotype (df = 2, p < 0.001; partial R^2^ = 0.59), but no significant effect of latitude (df = 1, p = 0.403; partial R^2^ = 0.03). Both Antarctic (types B1, B2 and C) and Atlantic Type 1 (Iceland and Norway) ecotypes had significantly higher sums of short ROH relative to ‘other’ ecotypes, despite an overlap in latitudinal range. The North Pacific *resident* ecotype, which was excluded from the model due to a low sample size, had a similarly high sum of short ROH as the Antarctic ecotypes (Figure 2D). In contrast, the sum of ROH greater than 2 Mb in each individual genome showed no significant relationship with latitude (Spearman correlation ρ = - 0.28, P = 0.17; Figure S6B). This result was supported by a GLM which also included ecotype as a predictor; neither predictors was significant (ecotype: df = 2, p = 0.151, latitude: df = 1, p = 0.09; Table S6).

Since homozygous tract length (*L*) declines exponentially as a function of recombination rate (*r*) and time (*t*), ROH due to background inbreeding during an ancestral bottleneck are expected to be shorter than ROH caused by recent inbreeding [15]. If *L* is approximated as 1/(*rt*) [17], then solving this equation provides the approximate coalescent time of the two homologous haplotypes within a given ROH class length. Assuming a range of constant mammalian recombination rates (0.649-1.554 cM/Mb) [18], we estimate that ROH less than 1 Mb in length correspond to haplotypes representing coalescence between 2 – 12 KY BP. In contrast, ROH longer than 2Mb are estimated to result from inbreeding during the past ∼ 2,000 years (Figure S7). This estimated timing of an ancestral bottleneck from short ROH (< 1 Mb) corresponds well with the demographic histories inferred by the Pairwise Sequentially Markovian Coalescent (PSMC) method [19] (Figure S8). Both analyses reflect increased coalescence during the Holocene in some high latitude populations and are comparable to patterns attributed to serial founder effects in Eurasian modern human populations [19, 20].

Our findings from a global dataset of genomes reinterpret previous inference based upon just two genomes; this earlier study interpreted this sparse data as evidence for a widespread genetic bottleneck across most of the species range during the Last Glacial Maximum (LGM) [21]. Our results are instead consistent with the scenario suggested by Hoelzel et al. [7], in which distinct genetic bottlenecks occurred in the ancestors of present-day high latitude populations, caused by serial founder effect during the colonisation of emerging habitat post-LGM. This interpretation is supported by observations of these same high latitude populations being the most differentiated based on allele frequencies [8, 9].

### Runs of Homozygosity Reflect Inbreeding

Individual genomic inbreeding coefficients estimated from ROH (F_ROH_), obtained as the total sum of ROH > 1 Mb divided by the total length of autosomal > 10 Mb in length [15], were correlated with our estimates of θ (Spearman correlation ρ = −0.6, P = 0.0013; Figure S9). A geographically widespread set of 22 of the 26 samples were considered inbred based on having estimated inbreeding coefficients (the proportion of the autosomes in homozygous tracts > 1 Mb) of F_ROH_ < (Table S1; Figure S10). This value corresponds to the mean kinship in the pedigree being that of a second cousin pairing [22], and is concordant with previous estimates of pairwise relatedness due to identity-by-descent (IBD), i.e. genetic identity due to a recent common ancestor in high latitude ecotypes [14]. Our finding of widespread inbreeding across the species range is consistent with the close kin structure and high social philopatry observed in most killer whale populations studied to date (Table S7). These F_ROH_ estimates surpass that reported in other populations characterised as having long-term low *N*_e_, including bonobos and the Native American Karitiana of the Brazilian Amazon [20,23,24].

A killer whale from a small population studied off the West coast of Scotland was a clear outlier, with 41% of the autosomal regions comprised of ROH > 1 Mb (Figure S10). The distinct distribution of ROH lengths in this individual (Figure 2) indicated long-term inbreeding throughout the Holocene. This estimated inbreeding coefficient (F_ROH_) exceeds those estimated for the Altai Neanderthal [25], and eastern lowland and mountain gorillas [23,26], each of which are benchmark taxa for the impact of long-term inbreeding on heterozygosity. The inbreeding coefficient of the Scottish killer whale was comparable to those estimated for the iconic inbred populations of the grey wolves (*Canis lupus*) of the Isle Royale [27] and Scandinavia [28]. Long-term monitoring of this Scottish population using photo-identification data estimated an extremely small census size (< 10 individuals) and zero fecundity during over two decades of data collection [29]. As previously argued, the low adult mortality rate in this small population suggests that bioaccumulation of organic pollutants, prey depletion and lethal interactions with fisheries are not the principle drivers of the small size of this population [29]. However, lack of recruitment through reproduction suggests that the population has reached a critical point whereby it experiences inverse density dependence (the ‘Allee’ effect) due to a loss of fitness from inbreeding, demographic stochasticity and reduced benefits of sociality [29].

### Admixture reduces ROH at Low Latitudes

Genomes sampled at low latitudes, where killer whales are found at low density, were typically characterised as less autozygous than those sampled at high latitudes (Figure 2D; Table S1). Admixture between lineages with divergent evolutionary histories, and therefore distinct genetic backgrounds, is predicted to break up ROH, through the recombination of different haplotypes [15]. For example, comparison of patterns of ROH in a global dataset of human genomes found that African Americans had notably shorter mean ROH length than most Europeans or Africans [20]. We illustrate this outcome of episodic admixture of different genetic backgrounds by comparing the ROH of F1 offspring from Icelandic and North Pacific *transient* parents, with the ROH of the parental lineages (Figure 3, Figure S11). The two admixed individuals show a reduction in longer (> 1 Mb) ROH relative to the parental lineages, with recombination in subsequent generations expected to further break down shorter ROH. The relative genetic homogeneity among low latitude samples [8,9] suggests gene flow. However, explicit testing of admixture is needed. That we see fewer ROH in genomes from low latitude regions, where killer whale density is lowest [4], would suggest that episodic admixture between divergent lineages, (rather than continuous equilibrium gene flow), is the primary process breaking up ROH in these low latitude populations. Episodic admixture, as opposed to continuous equilibrium gene flow, is corroborated by the presence of some of the longest ROH (> 8 Mb) being found in low latitude samples, such as that from Clipperton Island in the Eastern Tropical Pacific (Figure 4), indicative of recent inbreeding events (estimated to have occurred in the past 10 generations, Figure S7).

**Figure 3.**
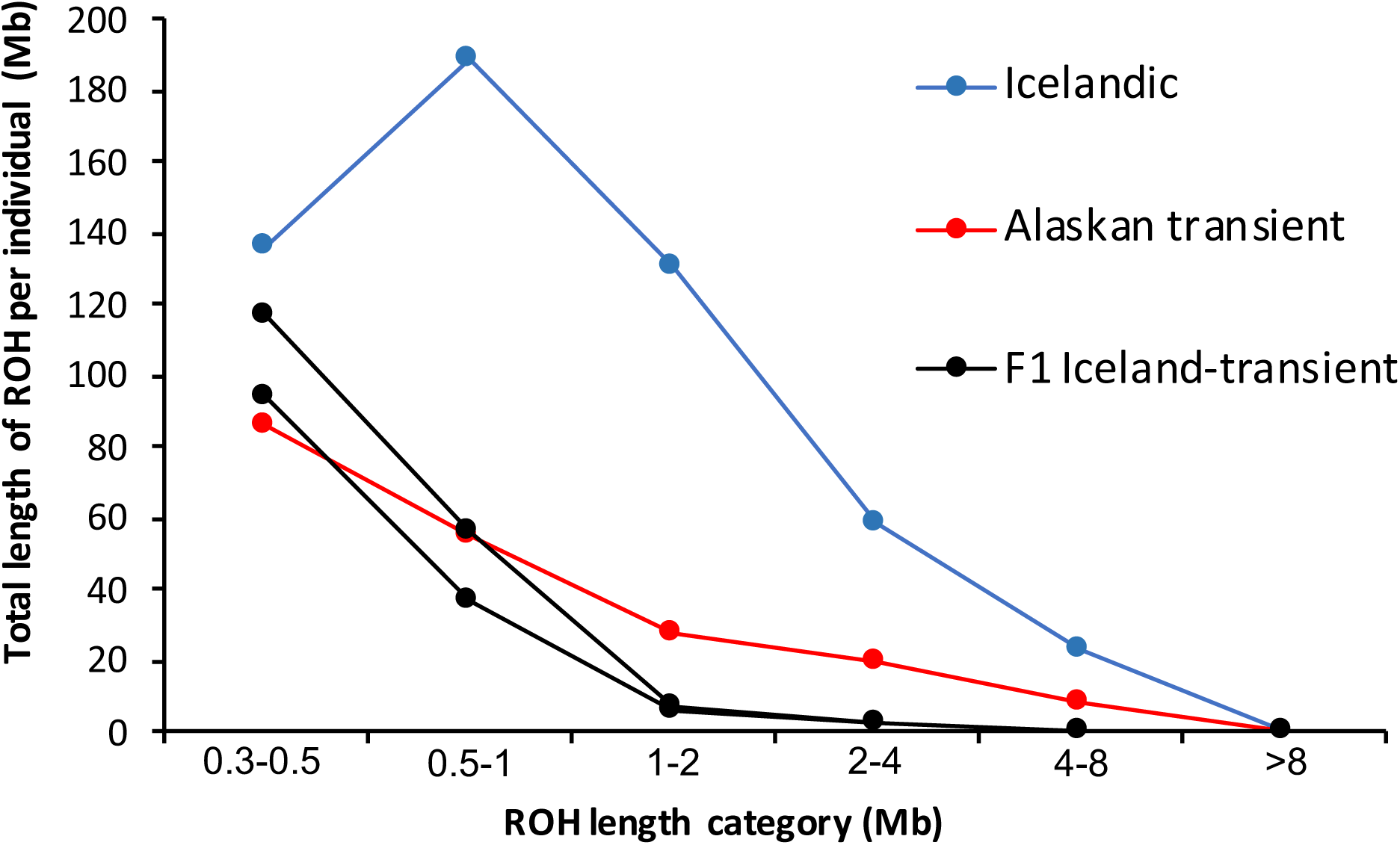
Comparison of ROH in F1 admixed individuals and their parental lineages. Sum of runs of homozygosity in different length categories in the genomes of two F1 admixed individuals and their North Atlantic and North Pacific parental lineages.

**Figure 4.**
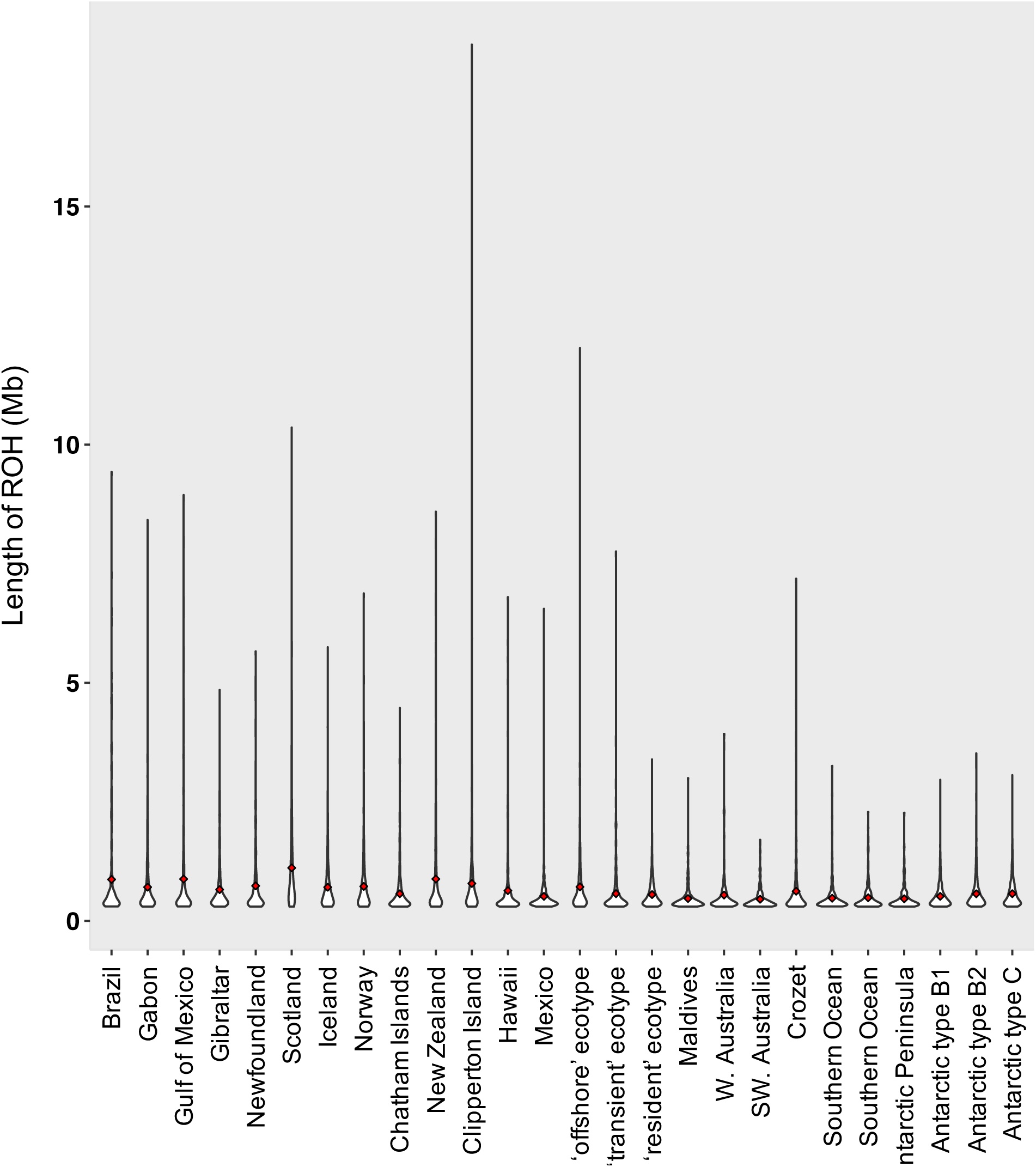
Kernel density plots of the length of runs of homozygosity (ROH) in individual genomes. Red markers indicate mean ROH length per individual. Samples are ordered approximately by sampling location (Atlantic, Pacific, Indian and Southern Ocean; and low to high latitude). The *offshore, transient* and *resident* ecotypes were all sampled in Alaska.

## DISCUSSION

Killer whales have radiated globally in a timescale comparable to anatomically modern humans. In this study, we have shown that like human populations that expanded out of Africa [15, 20], present day killer whale populations at the edge of the species’ range also share the genomic footprints of past demographic history, namely increased inbreeding and consequentially reduced heterozygosity.

Latitude was a predictor of patterns of heterozygosity, though the genomes of some high latitude populations did not have heterozygosity characteristic of a post-glacial expansion. As an example, in contrast to the North Pacific *resident* fish-eating ecotype, the sympatric *transient* mammal-eating ecotype had a similar pattern of heterozygosity and ROH to low-latitude populations. The *transient* ecotype was recently shown to be highly admixed, sharing partial ancestry with the *offshore* ecotype and low latitude populations [9]. Post-bottleneck admixture has been proposed to ‘rescue’ populations and reduce the recessive mutation load (dominance heterosis), in particular upon secondary contact after a period of allopatry and when the two admixing populations’ demographic histories do not include shared bottlenecks [30-33]. To the best of our knowledge and following five decades of field observations and annual censuses of populations [34], there is no evidence of greater fitness in the *transient* ecotype compared with the sympatric *resident* ecotype, despite the differences in genetic diversity, and the relationship between individual heterozygosity and fitness inferred from these long-term pedigree studies is equivocal [35]. The mutation load from post-bottleneck expansion (expansion load) is expected to comprise of many weakly deleterious recessive mutations in homozygous genotypes, which may have only a small effect on fitness [36], making it difficult to disentangle their impact from extrinsic influences on fitness. Additionally, if ROH are stable over time, then deleterious recessive mutations with homozygous genotypes become exposed to selection and can be purged over time [37, 38].

However, the potential fitness impacts of inbreeding are evident in the small (< 10 individuals) Scottish West Coast population, which has had zero recruitment over the past two decades [29]. The reduction in genetic diversity and individual fitness caused by inbreeding depression can impact population growth and increase the chance of population extinction [39, 40]. Our results suggest the extreme level of long-term inbreeding detected in the Scottish sample exceeds a threshold above which killer whale populations become unviable (see [29] for consideration of other threats versus inbreeding in this population). Consanguineous mating generates new recessive homozygotes, with an impact upon offspring fitness drawn from a distribution of fitness effects, which based upon human data includes an estimated equivalent of 3-5 lethal recessive mutations that act between late foetal and early adult stages [41]. Yet, even weakly deleterious recessive mutations can contribute to the heritability of complex traits such as genetic diseases and morphology [42]. Autosomal recessive disorders, for example Chédiak-Higashi syndrome [43], have been identified in wild killer whales, but remain under-studied. The genetic impacts of inbreeding depression should therefore be of key concern for populations already impacted by contaminants, depleted prey populations and lethal fishery interactions [29, 44-46].

## Methods

Our dataset consisted of twenty-six previously published genomes [9] from samples selected from a dataset of 452 individuals that best represent the known global geographic and genetic diversity of this species [8]. An additional two genomes from F1 admixed parentage of Icelandic mothers and a North Pacific *transient* father were generated as follows. Genomic DNA was sheared to an average size of ∼500 bp using a Diagenode Bioruptor Pico sonication device. The sheared DNA extracts were converted to blunt-end Illumina sequencing libraries using New England Biolabs (Ipswich, Ma., USA) NEBNext library kit E6040L [47]. Libraries were subsequently dual indexed and amplified for 15 cycles using a KAPA HiFi HotStart PCR kit (Kapa Biosystems, Wilmington, Ma. USA) in 50-µl reactions following the manufacturer’s guidelines. The amplified libraries were purified using a QIAquick PCR purification kit (Qiagen, Hilden, Germany) and size-selected on a 2% agarose gel in the range 422-580 bp using a BluePippin instrument (Sage Science, Beverly, Ma. USA). The DNA concentration of the libraries was measured using a 2100 Bioanalyzer (Agilent Technologies, CA, USA); these were then equimolarly pooled and sequenced across a lane of an Illumina HiSeq4000 platform. Read trimming, mapping, filtering and repeat-masking was conducted as per reference [9]. Briefly, reads from each individual were processed with AdapterRemoval2 [48] to trim residual adapter sequence contamination and to remove adapter dimer sequences as well as low-quality stretches at the read ends. Filtered reads >30 bp were then mapped using BWA-MEM algorithm to a high-quality, 2.25-Gb reference assembly (Oorca1.1) [49], requiring a mapping quality greater than 30. Clonal reads were collapsed using the rmdup function of the SAMtools [50]. Repetitive elements identified using RepeatMasker [51] and the Cetartiodactyl repeat library from Repbase [52], in addition to regions of low (less than a third of the mean) and excessive (more than twice the mean) coverage; regions of poor mapping quality (Q<30); and regions called as Ns in the reference sequence, were masked using BEDtools [53]. Sites were further filtered to include only autosomal regions.

The population mutation rate, theta (θ), which is equal to 4*N*_e_μ, was estimated directly from shotgun sequencing data mapped to the autosomal scaffolds >10Mb using the maximum likelihood estimator [54] implemented in mlRho v.2.9. [55]. The autosomal mutation rate (μ) was assumed to be 2.34×10^−8^ substitutions per nucleotide per generation [56]. We first estimated θ for 20× and randomly down-sampled to 5× coverage autosomal scaffolds of the Norwegian and resident killer whale genomes to establish a correction factor for our 5× genomes. We found that θ estimates from the 5× coverage genomes were consistently downwardly biased by a factor of approximately 0.75. We therefore report estimates of θ from our 5x coverage data after applying a correction factor of 1/0.75. To compare the relationship between measures of heterozygosity and latitude we used cor.test in R to calculate the Spearman correlation coefficient (ρ).

Runs of homozygous genotypes (ROH) were identified using the window-based approach implemented in PLINK v1.07 [16] from an input file of genotype likelihoods generated from autosomal scaffolds >10Mb by ANGSD [57] with the following filtering settings: removing reads of poor mapping quality (MAPQ < 30), removing sites with low base quality scores (q < 20), calling only SNPs inferred with a likelihood ratio test (LRT) of P < 0.000001, discarding reads that did not map uniquely, adjusting q-scores around indels, adjusting minimum quality score to 50 for excessive mismatches, and discarding bad reads (flag >=256). Inferred SNPs were lightly pruned based on linkage disequilibrium (LD) r^2^ >0.9 using PLINK, which has been found to improve the accuracy of detecting autozygous ROHs [58]. We estimated ROH from pruned and unpruned data and found minimal qualitative difference with our data. Sliding window size was set to 300 kb, with a minimum of 50 SNPs at a minimum density of 1 SNP per 50 kb required to call a ROH. To account for genotyping errors, we allowed up to 5 heterozygote sites per 300 kb window within called ROHs, as per ref. [59]. A length of 1,000 kb between two SNPs was required in order them to be considered in two different ROHs.

General linear models to test the influence of latitude and ecotype on θ, heterozygosity and the sum of short and long ROH were run using the package lme4 [60]in R-3.6.1 [61]. Ecotype can be a somewhat subjective categorisation [62]. We designated samples to the well-defined North Pacific *offshore, resident* and *transient* ecotypes [34], to the well-defined Antarctic morphotypes type B1, type B2 and type C [63], for which there is increasing evidence of correlated ecological variation [64,65], and to Atlantic type 1 which feeds mainly upon Atlantic herring, with some groups seasonally feeding upon seals [66]. These ecotype definitions are based upon ecological data, but are also reflected in the genetic data, representing distinct genetic clusters [8,9,14] All other individuals were classified as ‘other’, which is an under-estimation of the ecological variation or the degree of ecological specialisation among these samples, however, it does reflect the more homogenous genetic variation among these samples [8,9]. For θ and ROH models, ecotypes with one sample were removed (resident, transient and offshore). For the θ model, model plots indicated that two samples were resulting in a problematic model fit. These two samples (Scotland and Maldives) were therefore excluded. The removal of these samples did not substantially change model results. The Scottish sample was also removed from the long ROH model due to violations of model fit, but again, this did not have a substantial effect on model results. For the short ROH model, the response variable was log transformed to improve model fit. For each final model, model plots showed the assumptions of homogeneity and normality of residuals were met.

Habitat maps modelling the distribution of suitable habitat for killer whales at the present and during the LGM were originally constructed for a study by Morin et al. [8] using the AquaMaps approach [66,67]. Predictions of the relative environmental suitability for killer whales were projected into geographic space by relating habitat preferences (based on a predefined set of environmental parameters including depth, temperature, salinity and sea ice concentration) to local conditions using environmental data for different time periods and assuming no changes in species-specific habitat usage over time.

## Acknowledgements

The sequencing service was provided by the Genomics Core Facility (GCF), Norwegian University of Science and Technology (NTNU). GCF is funded by the Faculty of Medicine and Health Sciences at NTNU and Central Norway Regional Health Authority. A.D.F. was supported by a short visit grant from the European Science Foundation–Research Networking Programme ConGenOmics and by a Swiss National Science Foundation grant (31003A-143393) to L. Excoffier, and by the European Union’s Horizon 2020 research and innovation programme under the Marie Skłodowska-Curie grant agreement No. 663830. MTPG acknowledges ERC Consolidator Award 681396-Extinction Genomics. We also thank James McBain and Debbie Duffield for providing DNA.

