## Supplementary materials for "Runs of homozygosity in killer whale genomes provide a global record of demographic histories"

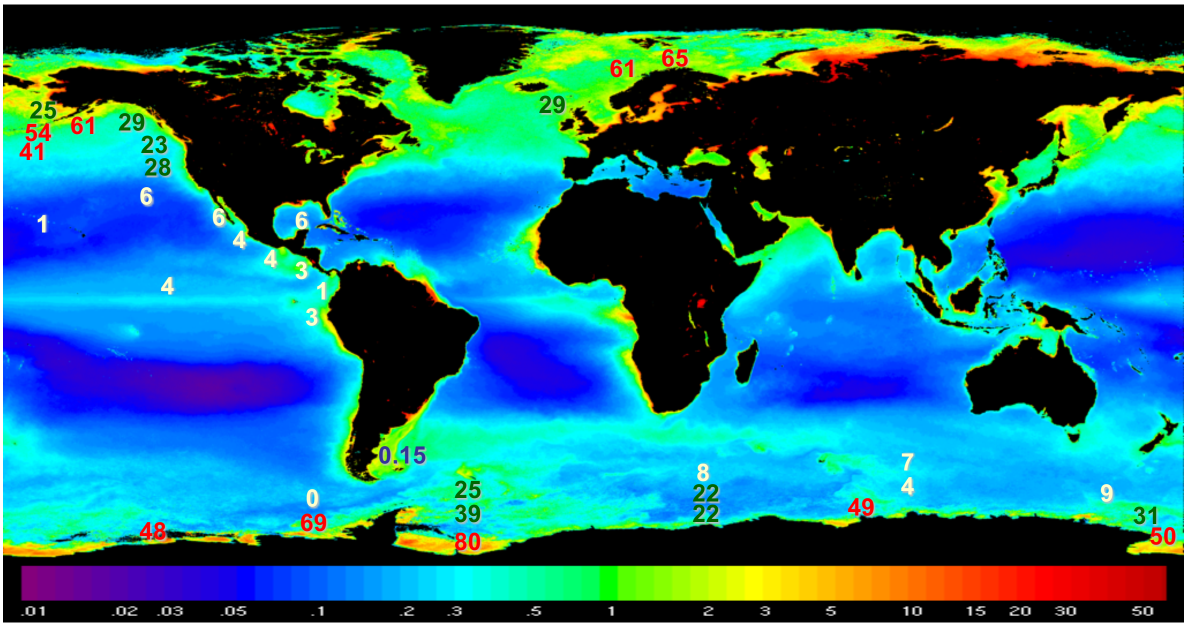


**Figure S1: World-wide killer whale densities.**
Numbers on map indicate killer whale density in animals per 10,000 $km^2$. Shading indicates ocean productivity as measured by the average chlorophyll-a concentration ($mg/m^3$) from 1997-2002 SeaWiFS images. Adapted from Forney & Wade [6].

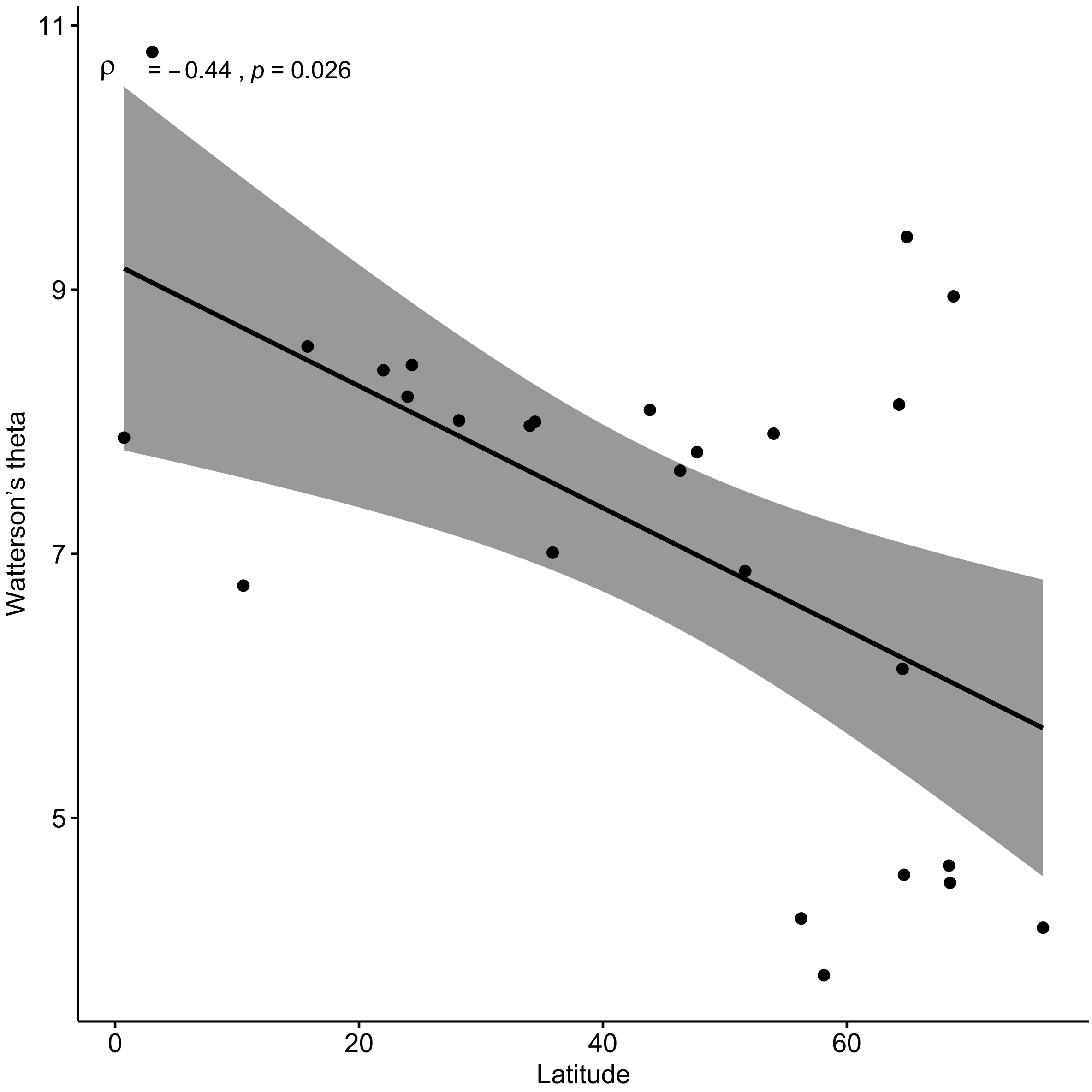


**Figure S2: Spearman Rank Correlation of Watterson's theta ($\theta$) with degrees latitude from the equator.** Shading indicates the 95% confidence intervals of the forecasted regression line.

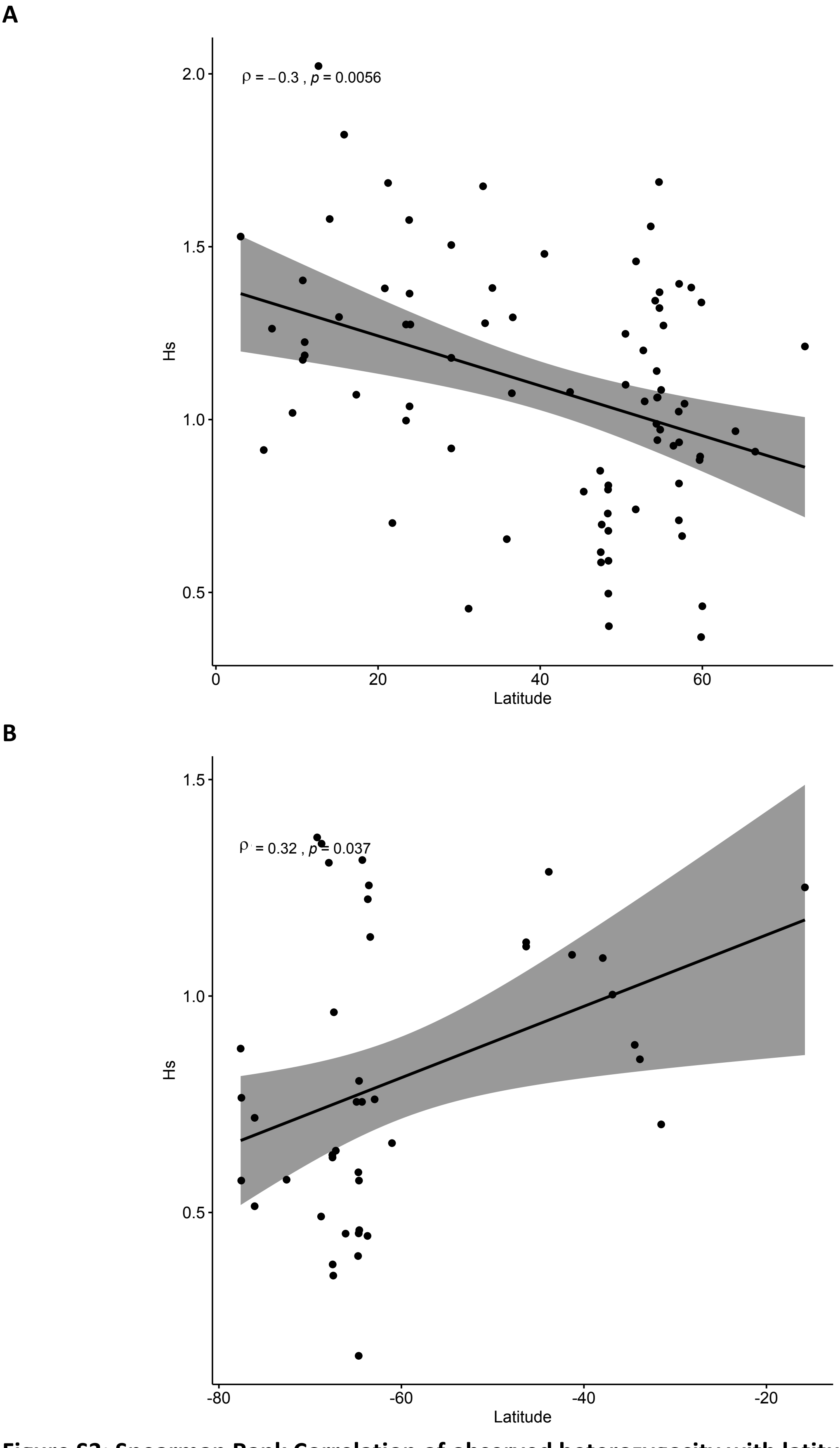


**Figure S3: Spearman Rank Correlation of observed heterozygosity with latitude.**
**A** Northern hemisphere and **B** Southern hemisphere. Shading indicates the 95% confidence intervals of the forecasted regression line.

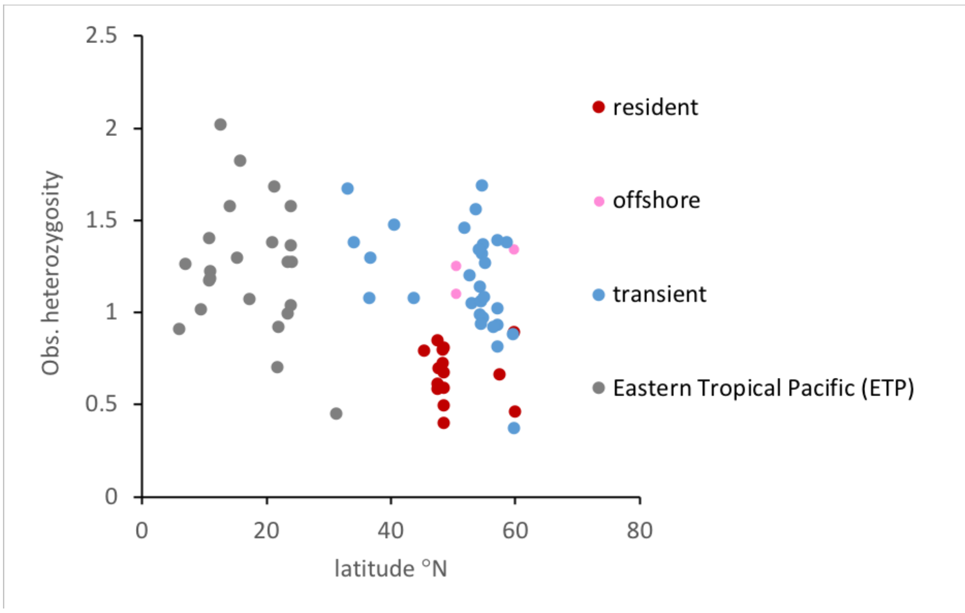


**Figure S4:** The relationship between latitude (*x*-axis), heterozygosity (*y*-axis) and ecotype (see key), based on genotypes of 71 individuals from the North Pacific taken from Morin et al. 2015 [8]. Ecotype explains more variance in heterozygosity than latitude (partial R-squared for ecotype = 0.37; partial R-squared for latitude = 0.09). Model summary is given in Table S3.

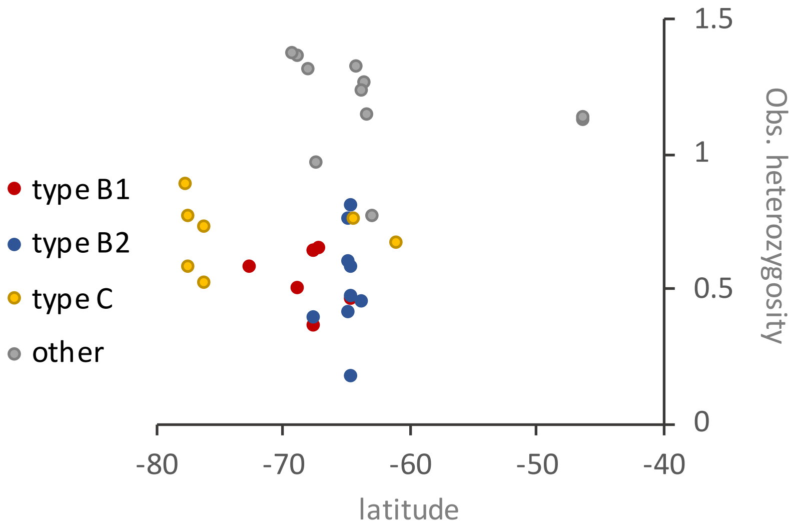


**Figure S5:** The relationship between latitude (*x*-axis), heterozygosity (*y*-axis) and ecotype (see key), based on genotypes of 35 individuals from the Antarctic and Southern Ocean taken from Morin et al. 2015 [8]. Ecotype explains more variance in heterozygosity than latitude (partial R-squared for ecotype = 0.77; partial R-squared for latitude = 0.02). Model summary is given in Table S4.

**A**

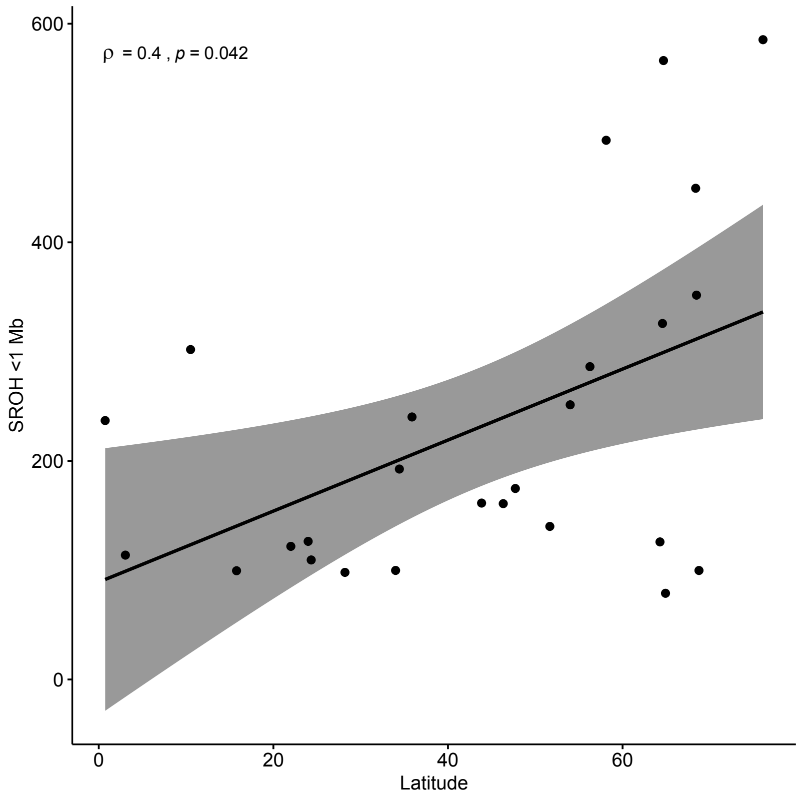


**B**

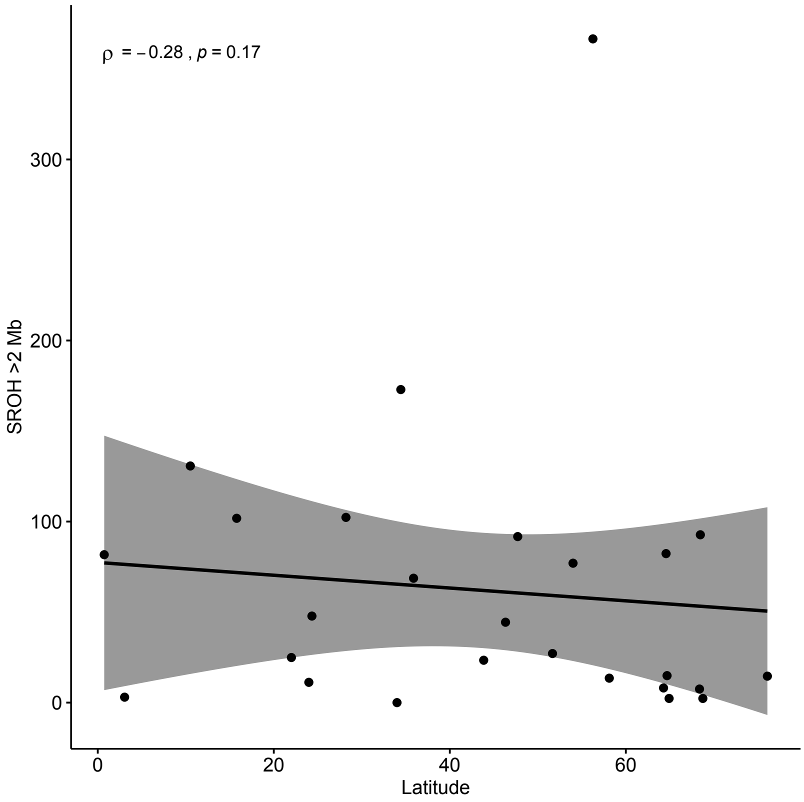


**Figure S6: Spearman Rank Correlation of the sum of runs of homozygosity (SROH) with latitude. A** ROH less than 1Mb in length and **B** ROH greater than 2 Mb in length. Units of the *y*-axes are Mb. Shading indicates the 95% confidence intervals of the forecasted regression line.

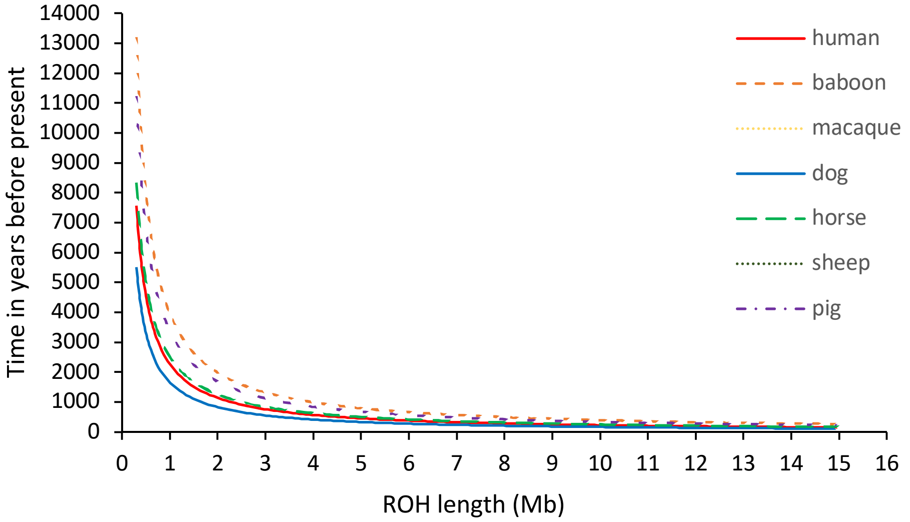


**Figure S7: Expected age of homozygous tracts,** assuming *L* (tract length) is approximated as 1/(*rt*) [17], and a generation time of 25.7 years [68] and a range of mammalian recombination rates (*r*) (as per [18]). If these assumptions are realistic then the ROH categories considered in this study would primarily reflect demographic processes during the Holocene.

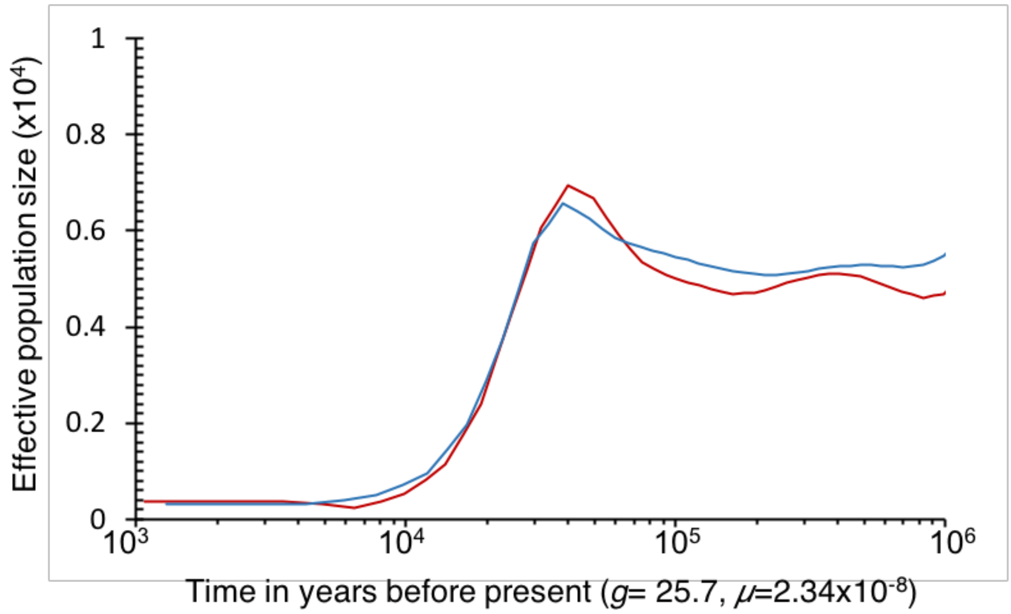


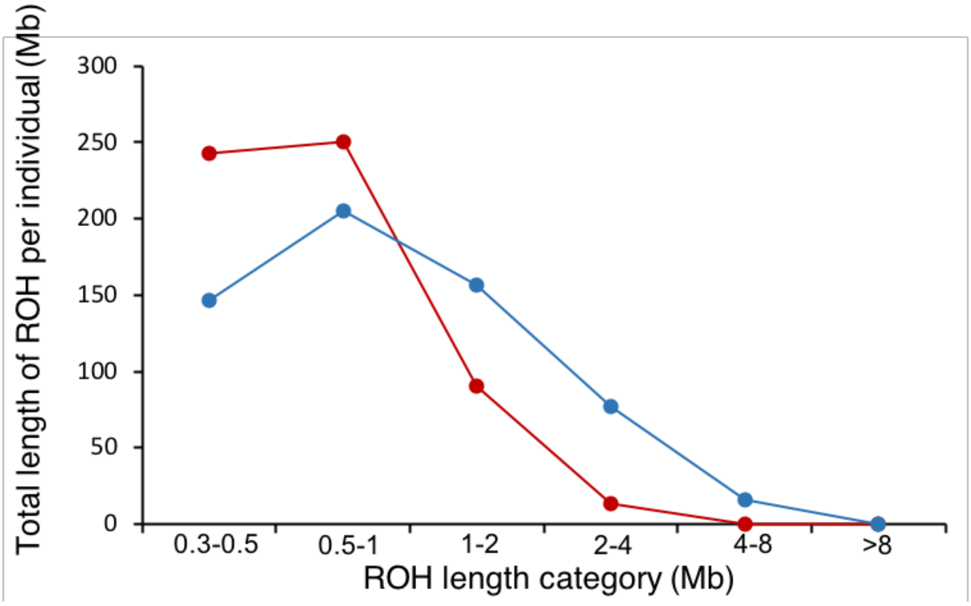


**Figure S8: Comparison of inferred effective population size through time and relative size of runs of homozygosity (ROH).** Plots represent data from the Alaskan resident (red) and Norway (blue) populations. Effective population size is inferred using the Pairwise sequentially Markovian coalescent (PSMC) as per [14]. The PSMC estimates of $N_e$ suggest a stable ancestral effective population size that is comparable to $N_e$ derived from many of the theta ($4N_e\mu$) estimates in Supplementary table 1 of $N_e$=0.5-1.0x10$^4$, and that likely represents the global effective population size. Both the resident and Norway populations are inferred by PSMC to have undergone extended genetic bottlenecks following the Last Glacial Maximum. PSMC infers $N_e$ through time from estimated changes in coalescent rates based on the frequency and length of homozygous tracts throughout the genome.

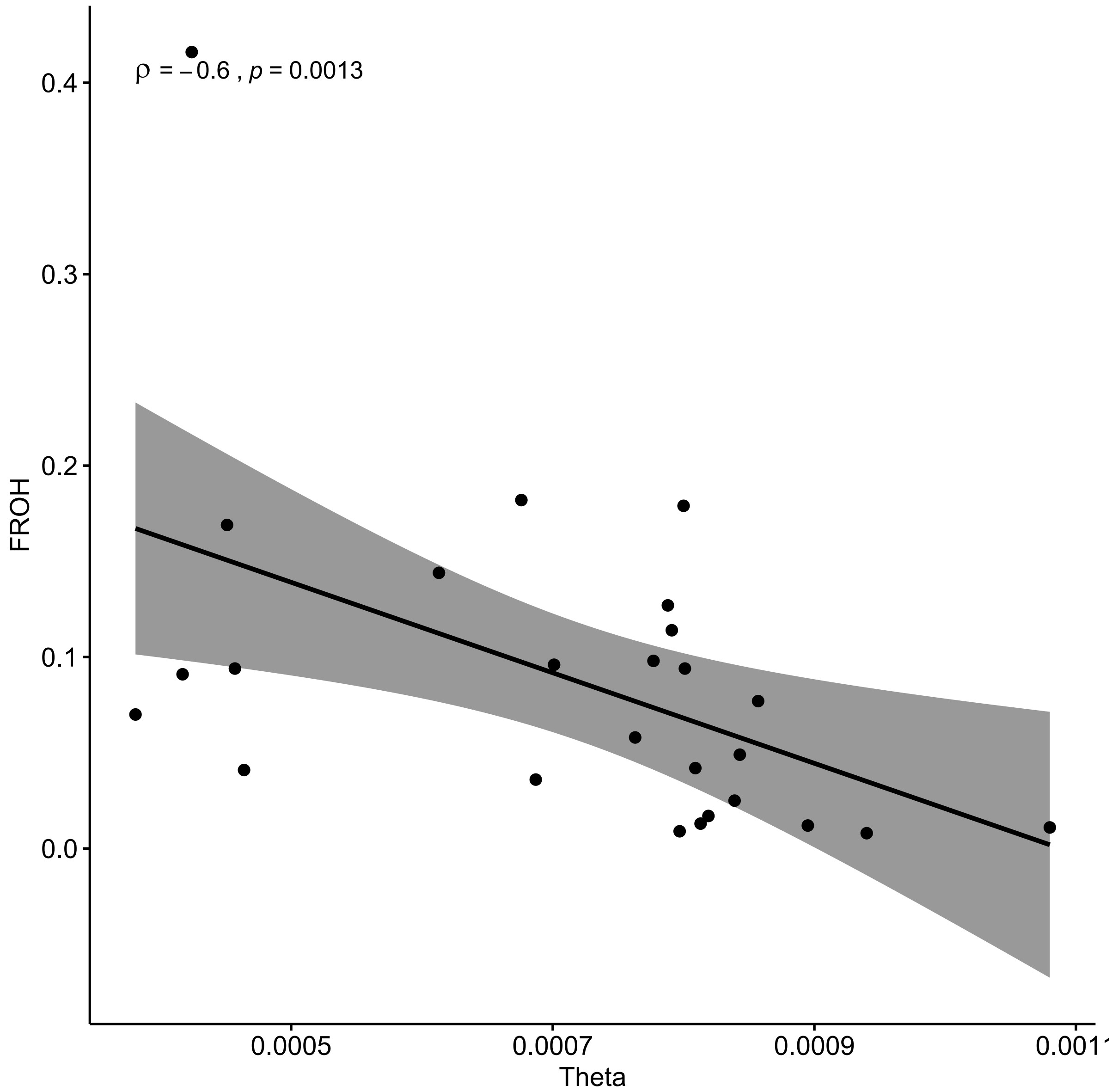


**Figure S9: Spearman Rank Correlation of theta with genomic inbreeding coefficient ($F_{ROH}$).** Shading indicates the 95% confidence intervals of the forecasted regression line.

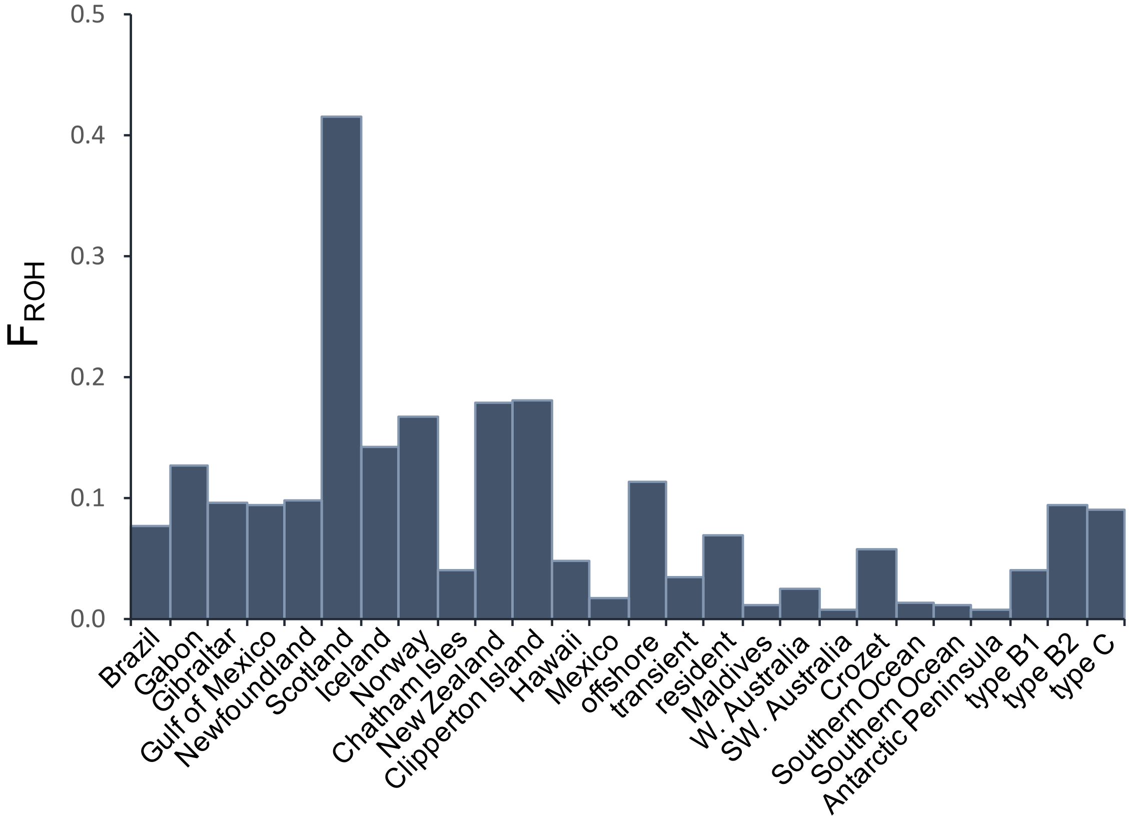


**Figure S10.** $F_{ROH}$ estimated as the proportion of the autosomal scaffolds which are >10Mb in length, which were covered by ROH $\geq$ 1Mb in length.

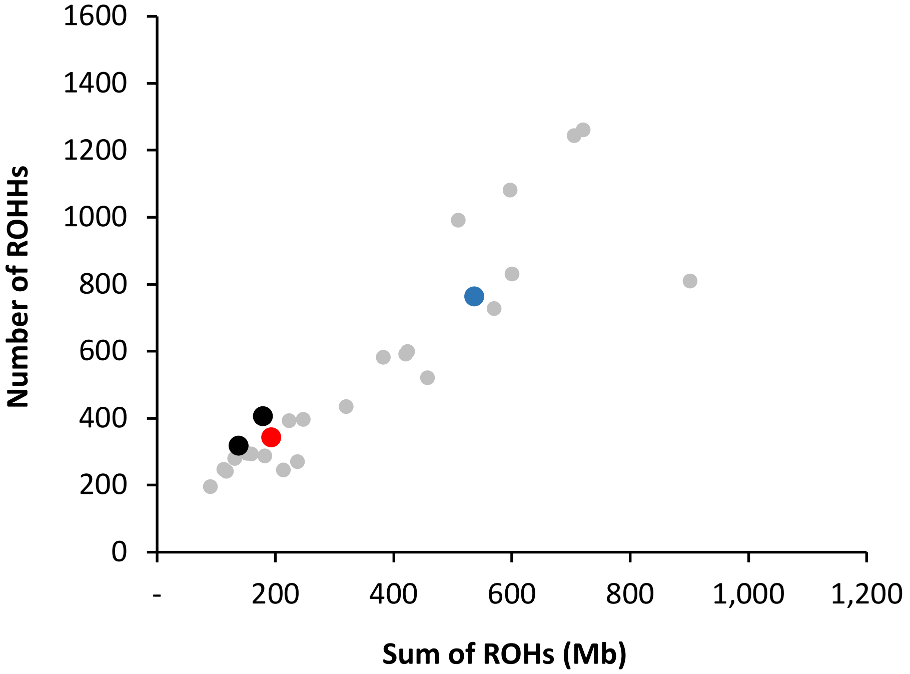


**Figure S11:** Number of ROHs compared to the total length of ROHs across the autosomes. The blue data point represents the Icelandic sample, the red data point the North Pacific transient sample and the two black data points represent the two F1 offspring of a North Pacific transient father and an Icelandic mother.

| Region | Population | *θ* (x$10^4$) | CI (x$10^4$) | $F_{ROH}$ |
|---|---|---|---|---|
| Pacific | Alaskan *transient* | 6.87 | 6.83 - 6.89 | 0.036 |
| | Alaskan *resident* | 3.81 | 3.79 - 3.84 | 0.070 |
| | Alaskan *offshore* | 7.91 | 7.87 - 7.95 | 0.114 |
| | Hawaii | 8.43 | 8.39 - 8.47 | 0.049 |
| | Clipperton Island (ETP) | 6.76 | 6.73 - 6.80 | 0.182 |
| | Mexico (ETP) | 8.19 | 8.15 - 8.21 | 0.017 |
| Atlantic | Norway | 4.51 | 4.48 - 4.53 | 0.169 |
| | Iceland | 6.13 | 6.11 - 6.16 | 0.144 |
| | Scotland | 4.24 | 4.21 - 4.27 | 0.416 |
| | Newfoundland | 7.77 | 7.73 - 7.80 | 0.098 |
| | Gibraltar | 7.01 | 6.99 - 7.05 | 0.096 |
| | Gulf of Mexico | 8.01 | 7.99 - 8.05 | 0.094 |
| | Gabon | 7.88 | 7.85 - 7.92 | 0.127 |
| | Brazil | 8.57 | 8.53 - 8.61 | 0.077 |
| Australasia | Chatham Islands | 8.09 | 8.05 - 8.13 | 0.042 |
| | New Zealand | 8.00 | 7.96 - 8.04 | 0.179 |
| | W. Australia | 8.39 | 8.35 - 8.43 | 0.025 |
| | SW. Australia | 7.97 | 7.93 - 8.01 | 0.009* |
| Antarctica & Southern Ocean | type B1 | 4.64 | 4.61 - 4.68 | 0.041 |
| | type B2 | 4.57 | 4.55 - 4.60 | 0.094 |
| | type C | 4.17 | 4.15 - 4.20 | 0.091 |
| | Crozet | 7.63 | 7.59 - 7.67 | 0.058 |
| | Antarctic Peninsula | 9.40 | 9.36 - 9.44 | 0.008* |
| | Southern Ocean | 8.13 | 8.09 - 8.17 | 0.013* |
| | Southern Ocean | 8.95 | 8.91 - 8.97 | 0.012* |
| Indian Ocean | Maldives | 10.8 | 10.7 - 10.8 | 0.011* |

**Table S1: Estimates of theta (θ ) and genomic inbreeding coefficients ($F_{ROH}$).** Inbreeding coefficients are calculated by dividing the total sum of ROH greater than 1Mb in scaffolds $\geq$10Mb by the total length of the autosomal regions in scaffolds $\geq$10Mb (1,477 Mb). Asterisks indicate non-inbred individuals, based upon $F_{ROH} < 0.015$, the value of inbreeding corresponding to the mean kinship of a second cousin pairing. All other individuals are considered inbred.

**Model: lm(Theta ~ Ecotype + Latitude)**

| Comparison | Variable | β | SE | t-value | p-value |
|---|---|---|---|---|---|
| **Antarctic** | **Ecotype (other)** | 4.06E-04 | 4.84E-05 | 8.382 | **<0.001** |
| | **Ecotype (type 1)** | 9.01E-05 | 5.73E-05 | 1.572 | 0.134 |
| **Other** | **Ecotype (type 1)** | -3.15E-04 | 5.35E-05 | -5.901 | **<0.001** |
| | **Latitude** | 1.28E-06 | 8.15E-07 | 1.575 | 0.134 |

| | df | SS | RSS | p-value |
|---|---|---|---|---|
| **Ecotype** | 2 | 3.12E-07 | 3.80E-07 | **<0.001** |
| **Latitude** | 1 | 9.76E-09 | 7.66E-08 | 0.09 |

| **Partial R-squared** | |
|---|---|
| **Ecotype** | 0.82 |
| **Latitude** | 0.12 |

**Table S2**: **GLM of investigating factors that explain variation in theta (θ).**
Top – summary of model results with every pairwise combination of ecotypes shown; middle – model results as summarised by drop1 (test = $\chi^2$); bottom – partial $R^2$ results. Resident, transient and offshore samples were removed due to a sample size of 1 per ecotype. Model plots revealed that residuals from the Scotland and Maldives samples (ecotype = other) deviated substantially from the normal distribution. These samples had a large influence on model results (Cook's Distance > 4/n), and were thus removed.

**Model: lm(Heterozygosity ~ Ecotype + Latitude)**

| Comparison | Variable | β | SE | t-value | p-value |
|---|---|---|---|---|---|
| **ETP** | **Ecotype (offshore)** | 0.428 | 0.243 | 1.762 | 0.083 |
| | **Ecotype (resident)** | -0.173 | 0.183 | -0.947 | 0.347 |
| | **Ecotype (transient)** | 0.351 | 0.184 | 1.908 | 0.061 |
| **Offshore** | **Ecotype (resident)** | -0.602 | 0.173 | -3.479 | **<0.001** |
| | **Ecotype (transient)** | -0.078 | 0.165 | -0.471 | 0.639 |
| **Resident** | **Ecotype (transient)** | 0.524 | 0.086 | 6.062 | **<0.001** |
| | **Latitude** | -0.012 | 0.005 | -2.516 | **0.01** |

| | df | SS | RSS | p-value |
|---|---|---|---|---|
| **Ecotype** | 3 | 2.906 | 7.796 | **<0.001** |
| **Latitude** | 1 | 0.469 | 5.359 | **0.01** |

| **Partial R-squared** | |
|---|---|
| **Ecotype** | 0.373 |
| **Latitude** | 0.087 |

**Table S3**: **GLM of investigating factors that explain variation in heterozygosity among North Pacific samples.**
Top – summary of model results with every pairwise combination of ecotypes shown; middle – model results as summarised by drop1 (test = $\chi^2$); bottom – partial $R^2$ results

**Model: lm(Heterozygosity ~ Ecotype + Latitude)**

| Comparison | Variable | β | SE | t-value | p-value |
|---|---|---|---|---|---|
| **B1** | **Ecotype (B2)** | -0.018 | 0.085 | -0.21 | 0.835 |
| | **Ecotype (C)** | 0.136 | 0.092 | 1.472 | 0.152 |
| | **Ecotype (other)** | 0.659 | 0.086 | 7.703 | **<0.001** |
| **B2** | **Ecotype (C)** | 0.153 | 0.093 | 1.647 | 0.11 |
| | **Ecotype (other)** | 0.677 | 0.08 | 8.935 | **<0.001** |
| **C** | **Ecotype (other)** | 0.523 | 0.01 | 5.364 | **<0.001** |
| | **Latitude** | -0.004 | 0.005 | -0.792 | 0.435 |

| | df | SS | RSS | p-value |
|---|---|---|---|---|
| **Ecotype** | 3 | 2.651 | 3.445 | **<0.001** |
| **Latitude** | 1 | 0.017 | 0.811 | 0.394 |

| **Partial R-squared** | |
|---|---|
| **Ecotype** | 0.769 |
| **Latitude** | 0.021 |

**Table S4**: **GLM of investigating factors that explain variation in heterozygosity among Southern Ocean/Antarctic samples.**

Top – summary of model results with every pairwise combination of ecotypes shown; middle – model results as summarised by drop1 (test = $\chi^2$); bottom – partial $R^2$ results

**Model: lm(log(sumROH < 1mb) ~ Ecotype + Latitude)**

| Comparison | Variable | β | SE | t-value | p-value |
|---|---|---|---|---|---|
| Antarctic | Ecotype (other) | -1.416 | 0.283 | -5 | **<0.001** |
| | Ecotype (type 1) | -0.46 | 0.347 | -1.323 | 0.202 |
| Other | Ecotype (type 1) | 0.956 | 0.316 | 3.022 | **0.007** |
| | Latitude | -0.003 | 0.004 | -0.766 | 0.453 |

| | df | SS | RSS | p-value |
|---|---|---|---|---|
| **Ecotype** | 2 | 3.916 | 6.66 | **<0.001** |
| **Latitude** | 1 | 0.085 | 2.832 | 0.403 |

| **Partial R-squared** | |
|---|---|
| **Ecotype** | 0.588 |
| **Latitude** | 0.03 |

**Table S5**: **GLM of investigating factors that explain variation in the sum of short (<1Mb) ROH.** Top – summary of model results with every pairwise combination of ecotypes shown; middle – model results as summarised by drop1 (test = $\chi^2$); bottom – partial $R^2$ results. The response variable was log transformed in order to improve model fit. Resident, transient and offshore samples were removed due to a sample size of 1 per ecotype.

**Model: lm((sumROH > 2mb) ~ Ecotype + Latitude)**

| Comparison | Variable | β | SE | t-value | p-value |
|---|---|---|---|---|---|
| Antarctic | Ecotype (other) | 0.956 | 35.325 | 0.282 | 0.781 |
| | Ecotype (type 1) | 72.419 | 42.294 | 1.712 | 0.104 |
| Other | Ecotype (type 1) | 62.462 | 39.196 | 1.594 | 0.128 |
| | Latitude | -0.874 | 0.557 | -1.57 | 0.134 |

| | df | SS | RSS | p-value |
|---|---|---|---|---|
| **Ecotype** | 2 | 7239 | 45809 | 0.151 |
| **Latitude** | 1 | 5281.2 | 43851 | 0.093 |

**Table S6**: **GLM of investigating factors that explain variation in the sum of long (>2Mb) ROH.** Top – summary of model results with every pairwise combination of ecotypes shown; bottom – model results as summarised by drop1 (test = $\chi^2$). Resident, transient and offshore samples were removed due to a sample size of 1 per ecotype. Model plots revealed that residuals from the Scottish sample (ecotype = other) deviated substantially from the normal distribution, and that this sample had a large influence on model results (Cook's Distance > 4/n). This sample was thus removed.

| Ocean Basin | Population(s) | References |
|---|---|---|
| North Pacific | *resident* fish-eating ecotype | 69, 70, 71, 72, 73, 75 |
| | *transient* mammal-eating ecotype | 70, 74, 75 |
| | Hawaii | 76 |
| North Atlantic | Iceland/Scotland | 77,78,79 |
| Southern Ocean | Crozet and Marion Island | 80, 81 |

**Table S7**: Populations in which close kin-structure and social philopatry have been reported.
